# Glucose-stimulated calcium dynamics in beta cells from C57BL/6J, C57BL/6N, and NMRI mice: A systematic comparison of activation, activity, and deactivation properties in tissue slices

**DOI:** 10.1101/2022.01.14.476318

**Authors:** Viljem Pohorec, Lidija Križančić Bombek, Maša Skelin Klemen, Jurij Dolenšek, Andraž Stožer

## Abstract

Although mice are a very instrumental model in islet beta cell research, possible phenotypic differences between strains and substrains are largely neglected in the scientific community. In this study, we show important phenotypic differences in beta cell responses to glucose between NMRI, C57BL/6J, and C57BL/6N mice, i.e., the three most commonly used strains. High-resolution multicellular confocal imaging of beta cells in acute pancreas tissue slices was used to measure and quantitatively compare the calcium dynamics in response to a wide range of glucose concentrations. Strain- and substrain-specific features were found in all three phases of beta cell responses to glucose: a shift in the dose-response curve characterizing the delay to activation and deactivation in response to stimulus onset and termination, respectively, and distinct concentration-encoding principles during the plateau phase in terms of frequency, duration, and active time changes with increasing glucose concentrations. Our results underline the significance of carefully choosing and reporting the strain to enable comparison and increase reproducibility, emphasize the importance of analyzing a number of different beta cell physiological parameters characterizing the response to glucose, and provide a valuable standard for future studies on beta cell calcium dynamics in health and disease.

## INTRODUCTION

Laboratory mice are a vital source of islets of Langerhans in research on beta cell physiology, functional adaptation, and dysfunction during development of type 2 diabetes mellitus (T2DM) (50, 73), mostly due to their low housing costs, short breeding interval, availability for genetic manipulation, and regulatory requirements. From the translational point of view, despite some important differences(7) the islets of Langerhans show many structural and functional similarities in mice and men (22, 78). Therefore, it is not surprising that mouse models of T2DM exhibit comparable disease characteristics and can provide a significant insight into the mechanisms of T2DM development in humans (50). However, the translational relevance (16) is not the only important aspect when it comes to mouse models of beta cells physiology and pathophysiology. Another very important aspect are possible differences between individual mouse models (4). More specifically, either genetically defined inbred strains (27) or genetically undefined outbred stocks (19) are available. Some advocate greater use of inbred strains, arguing that the reduced genetic variability and the concomitantly reduced phenotypic variation contribute to the power of experimental results (19, 26–28). However, there is still uncertainty as to whether this is necessarily preferable, as comparable phenotypic variation has been demonstrated in inbred and outbred mice (13, 51, 88). Adding to the complexity is the fact that even substrains of the inbred strains can exhibit considerable phenotypic variation. The two most commonly used substrains – C57BL/6J and C57BL/6N, while descended from the parent strain C57BL/6, demonstrate genetic (64, 77) and phenotypic (61, 77) differences. One of the more pertinent differences in this regard is the nicotinamide nucleotide transhydrogenase (*Nnt*) gene, which encodes a mitochondrial enzyme involved in NADPH production (9) that harbors a mutation in the C57BL/6J substrain (31, 48, 87), whereas it remains intact in several C57BL/6N substrains (64). More specifically, at the phenotype level, examination of glucose handling in the two substrains yielded conflicting results. The NNT activity defect in C57BL/6J mice seems to be associated with attenuated glucose-stimulated insulin release (55, 87) compared to C57BL/6N mice (25, 49, 55, 87), while some report conflicting results for the two substrains (3, 25, 49, 92). Moreover, the two substrains are differentially sensitive to diet-induced obesity (DIO). Both C57BL/6J and C57BL/6N are prone to DIO, which is manifested by a marked increase in body mass (49, 65, 74), impaired glucose tolerance (29, 65, 74), an increase in fasting blood glucose and serum insulin that develops later in C57BL/6J than C57BL/6N (74), decreased insulin release (29), and insulin resistance (49). For a more detailed comparison of DIO phenotypes in different C57BL/6 substrains, see (49). In contrast, the outbred NNT competent NMRI mice exhibit better glucose handling compared with the C57BL/6 strain. The NMRI mice were reported to clear glucose more efficiently than the C57BL/6J (1), resulting in normal glucose tolerance (67). Moreover, they seem to be less susceptible to DIO because (i) feeding with HFD or WD resulted in a smaller increase in body mass than in C57BL/6J (1) or C57BL/6N (51) mice and (ii) glucose tolerance was less impaired than in C57BL/6J mice (1). Since DIO induced a comparable degree of insulin resistance in NMRI and C57BL/6J mice, normoglycemia was probably maintained in NMRI mice due to concomitant hyperinsulinemia (1). In the light of the above phenotypic differences among the mouse strains and substrains, one should be careful when generalizing findings on the molecular mechanisms obtained from a particular mouse strain (4).

For insulin-secreting beta cells, intracellular calcium concentration ([Ca^2+^]_i_) remains a suitable surrogate for the study of molecular machinery responsible for insulin secretion, and the importance of [Ca^2+^]_i_ dynamics for pancreatic insulin secretion is well acknowledged (35, 40, 41, 91). In general, stimulation of beta cells with glucose results in an initial transient increase in [Ca^2+^]_i_ in the first phase, followed by oscillatory changes in calcium influx in the second phase that persist until stimulation ceases (35, 36, 79). These changes in [Ca^2+^]_i_ are preceded by changes in membrane potential (23, 75) and drive insulin secretion (11, 37). Over the past century, numerous mouse models have been introduced to study [Ca^2+^]_i_ dynamics, most notably the genetically undefined outbred NMRI mice (36, 38, 54), and the *Ob/Ob* (*Lep*^ob^) mice (11, 39, 66) derived from a non-inbred colony in the middle of the previous century. However, the last two decades have witnessed greater diversity in the use of mouse models, often using genetically defined inbred strains, such as the C57BL/6J (59, 86) and the C57BL/6N mice (93), as well as genetically modified mice with the C57BL/6J background (17, 52, 90). We recently quantified the concentration-dependence of the beta cell response to glucose using high-resolution multicellular [Ca^2+^]_i_ imaging in many beta cells in tissue slices (85), but only for the NMRI strain. Given an increasing use of inbred strains and related knock-out models in islet research, it is important to assess whether the genetic background in general and the *Nnt* mutation in specific influence glucose-stimulated [Ca^2+^]_i_ dynamics in beta cells.

In this study, we therefore aimed to characterize and compare beta cell [Ca^2+^]_i_ dynamics in the three today most commonly used mouse strains, i.e., the NNT-deficient inbred C57BL/6J mice, NNT-competent inbred C57BL/6N mice, and NNT-competent outbred NMRI mice. To increase the probability of detecting possible changes, a wide range of physiological (7–10 mM) and supraphysiological (12-16 mM) glucose concentrations were used to stimulate beta cells in islets of Langerhans in acute pancreas tissue slices and their responses to glucose were systematically analyzed over the whole response, i.e., during activation, plateau activity, and deactivation.

## METHODS AND MATERIALS

### Ethics statement

The study protocol was approved by the Veterinary Administration of the Republic of Slovenia (approval number: U34401-12/2015/3 and U34401-35/2018-2). The study was conducted in strict accordance with all national and European recommendations pertaining to care and work with laboratory animals, and every effort was made to minimize animal suffering.

### Animals

Experiments were performed on 12–13-week-old male C57BL/6J, C57BL/6N, and NMRI mice, all acquired from Charles River. The mice were fed a standard rodent diet (Ssniff, Soest, Germany) and water *ad libitum*. They were housed in individually ventilated cages (Allentown LLC, USA) in groups of 1-4 animals per cage at 20-24 °C, 45-65 % relative humidity, and a 12-hour day-night lighting cycle.

Mice were weighed before sacrifice. Following the sacrifice with CO_2_ and cervical dislocation, the mouse was placed on its abdomen, and body length was measured from the nose tip to the base of the tail. Glucose was measured from the tail vein using the Accu-Check Aviva glucometer (Roche, Switzerland). During analysis, body mass index (BMI) was calculated as the ratio between body weight and body surface area (g/m2), where body surface area was calculated according to the following formula (33):

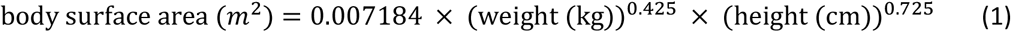

### Tissue slice preparation and dye loading

Acute pancreas tissue slices from C57BL/6J, C57BL/6N, and NMRI mice were prepared as described previously (80, 82, 83). In brief, after cervical dislocation of the animal, the abdominal cavity was accessed via laparotomy. The pancreas was injected with 1.9 % low-melting-point agarose (Lonza Rockland Inc., USA) at the proximal common bile duct, which was clamped distally at the major duodenal papilla. The agarose was dissolved in extracellular solution (ECS) containing (in mM 125 NaCl, 26 NaHCO_3_, 6 glucose, 6 lactic acid, 3 myo-inositol, 2.5 KCl, 2 Na-pyruvate, 2 CaCl_2_, 1.25 NaH_2_PO_4_, 1 MgCl_2_, 0.5 ascorbic acid) and maintained at 40 °C. Immediately following the agarose injection, the pancreas was cooled with the ice-cold ECS, extracted from the animal, and placed in a Petri dish containing cooled ECS. The tissue was then cut into 3-5 mm3 pieces, which were embedded in agarose and cut into 140 μm thick slices vibratome (VT 1000 S, Leica). The slices were transferred into a Petri dish containing HEPES-buffered saline (HBS, consisting of (in mM) 150 NaCl, 10 HEPES, 6 glucose, 5 KCl, 2 CaCl_2_, 1 MgCl_2_; titrated to pH=7.4 with 1 M NaOH) at room temperature. The prepared slices were stained in the dye-loading solution (6 μM Oregon Green 488 BAPTA-1 AM (OGB-1, Invitrogen), 0.03 % Pluronic F-127 (w/v), and 0.12 % dimethyl sulfoxide (v/v) dissolved in HBS) for 50 min at room temperature. All chemicals were obtained from Sigma-Aldrich (St. Louis, Missouri, USA) unless otherwise stated.

### Experimental protocol

Stained tissue slices were placed individually under the microscope into the recording chamber, which was continuously perifused with carbogenated ECS containing 6 mM glucose at 37 °C. To stimulate the slices, we manually changed the perifusate to a single stimulatory glucose concentration (7, 8, 9, 10, 12 or 16 mM) dissolved in carbogenated ECS at 37 °C for 40, 30, 20, 20, 15, and 15 min, respectively. After stimulation, the slice was reintroduced to the perifusate containing 6 mM glucose in ECS for at least 15 min.

### Imaging of intracellular free calcium concentration dynamics in beta cells

[Ca^2+^]_i_ imaging was performed using an upright confocal microscope system Leica TCS SP5 AOBS Tandem II with a 20X HCX APO L water immersion objective, NA 1.0, and an inverted confocal system Leica TCS SP5 DMI6000 CS with a 20X HC PL APO water/oil immersion objective, NA 0.7 (all from Leica Microsystems, Germany). The time series were acquired at a resolution of 512 × 512 pixels with a frequency of 2 Hz. The calcium reporter dye was excited with a 488 nm argon laser line, and the emitted fluorescence was detected with a Leica HyD hybrid detector in the 500-700 nm range (all from Leica Microsystems, Germany), as previously described (82, 83, 85).

### Data analyses

The cross-sectional areas of the islets were measured manually using high-resolution images (1024 × 1024 pixels) in the focal plane, where the [Ca^2+^]_i_ signals were recorded using LASX software (Leica Microsystems, Germany). For calcium dynamics analysis, regions of interest (ROI) were manually selected and exported using custom offline software (ImageFiltering, copyright Denis Špelič). Traces with considerable motion artifacts were excluded from further analysis, which was performed using in-house MATLAB scripts. A combination of linear and exponential fitting was used to account for the photobleaching effect. Activation times (i.e., the time delay from the onset of the stimulus to the transient increase in [Ca^2+^]_i_) and deactivation times (i.e., the time delay from the withdrawal of the stimulus to the last oscillation) for individual cells were determined manually using custom Matlab scripts. For the plateau phase of the beta cell responses, we binarized the oscillatory activity and defined the onset and termination of each oscillation at times of their half-maximal amplitudes using Matlab. The binarized signals were subsequently used to calculate the duration (i.e., the time interval between the onset and termination of a single oscillation), frequency (i.e., the inverse value of the interval between the onsets of two successive oscillations), and active time (i.e., the percentage of time occupied by oscillations). Active time (AT) is mathematically defined as the following product:

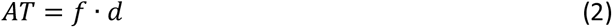

where *f* is the frequency (in Hz) and *d* is the duration (in seconds) of oscillations. Data were pooled across all cells and plotted using Tukey-style boxplots with whiskers denoting 1.5-times the interquartile range of the data. Statistical analyses were performed using GraphPad Prism 9.2 (GraphPad Software, Inc., San Diego, CA). Statistical differences between groups were tested using one-way analysis of variance (ANOVA) on ranks, followed by Dunn’s multiple comparison test, where data were not normally distributed, and one-way analysis of variance (ANOVA), where data were normally distributed. Asterisks denote statistically significant differences, as follows: * p<0.05, ** p<0.01, *** p<0.001, and **** p<0.0001. Shifts in the dose-dependent activation curves were calculated assuming a sigmoid-shaped 4 parameter logistic mathematical model (76), according to the following equation:

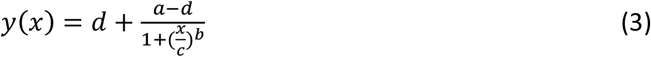

where delay in response *f(x)* is expressed as a function of concentration *x*, parameters *a* and *d* denote the lower and upper asymptote, respectively, *b* the slope of the linear portion of the curve, and *c* (commonly termed EC50/IC50) concentration at response midway between *a* and *d*. To compare the curves, parameters were set such that the value of the correlation coefficient was R^2^ ≥ 0.90 for the individual fits through median delays (*b* = −10.5, *a* = [100 200] s, and d = 850 s).

## RESULTS

In the study of the (patho)physiology of the pancreas, little attention is paid to differences between strains of mice. We therefore performed a systematic comparison between the three commonly used mouse strains, namely the C57BL/6J, C57BL/6N, and NMRI mice. We analyzed the beta cell (i) activation properties, (ii) oscillatory dynamics during the plateau phase of the response, and (iii) deactivation properties over a wide range of glucose concentrations. Additionally, we characterized the strains by using morphological measurements and by measuring non-fasting glucose.

### Gross morphological and glucose-handling interstrain differences

To describe the morphological differences between the strains, we measured body mass and nose-to-tail length. NMRI mice were larger (nose-to-tail length, median value 10.4 cm) and heavier (median value 42.9 g) than C57BL/6J (median length 8.9 cm and mass 24.8 g) and C57BL/6N mice (median length 9.0 cm and mass 27.5 g) mice (Figure 1 A&B), whereas the two inbred strains showed no difference in nose-to-tail length and body mass. Additionally, the calculated body mass index (BMI) was statistically significantly larger in NMRI mice than in either of the B6 strain (Figure 1C). Glucose handling, measured as non-fasting glucose levels, did not differ between strains (median values 7.6, 7.6, and 7.5 mM for C57BL/6J, C57BL/6N, and NMRI, respectively) (Figure 1 D). We also compared the gross morphology of the islets of Langerhans between C57BL/6J, C57BL/6N, and NMRI mouse strains (Figure 1E&F). Qualitatively, we found no differences in islet morphology between the three strains. To quantify the islet morphology, we measured the cross-sectional area of each islet in pancreas tissue slices that were subsequently designated for calcium imaging. Islets from the C57BL/6J mice were 33 % smaller by median compared with the NMRI mice and had a median of 10 % smaller cross-sectional area than the C57BL/6N mice, with no significant difference between the NMRI and C57BL/6N mice.

**Figure 1:**
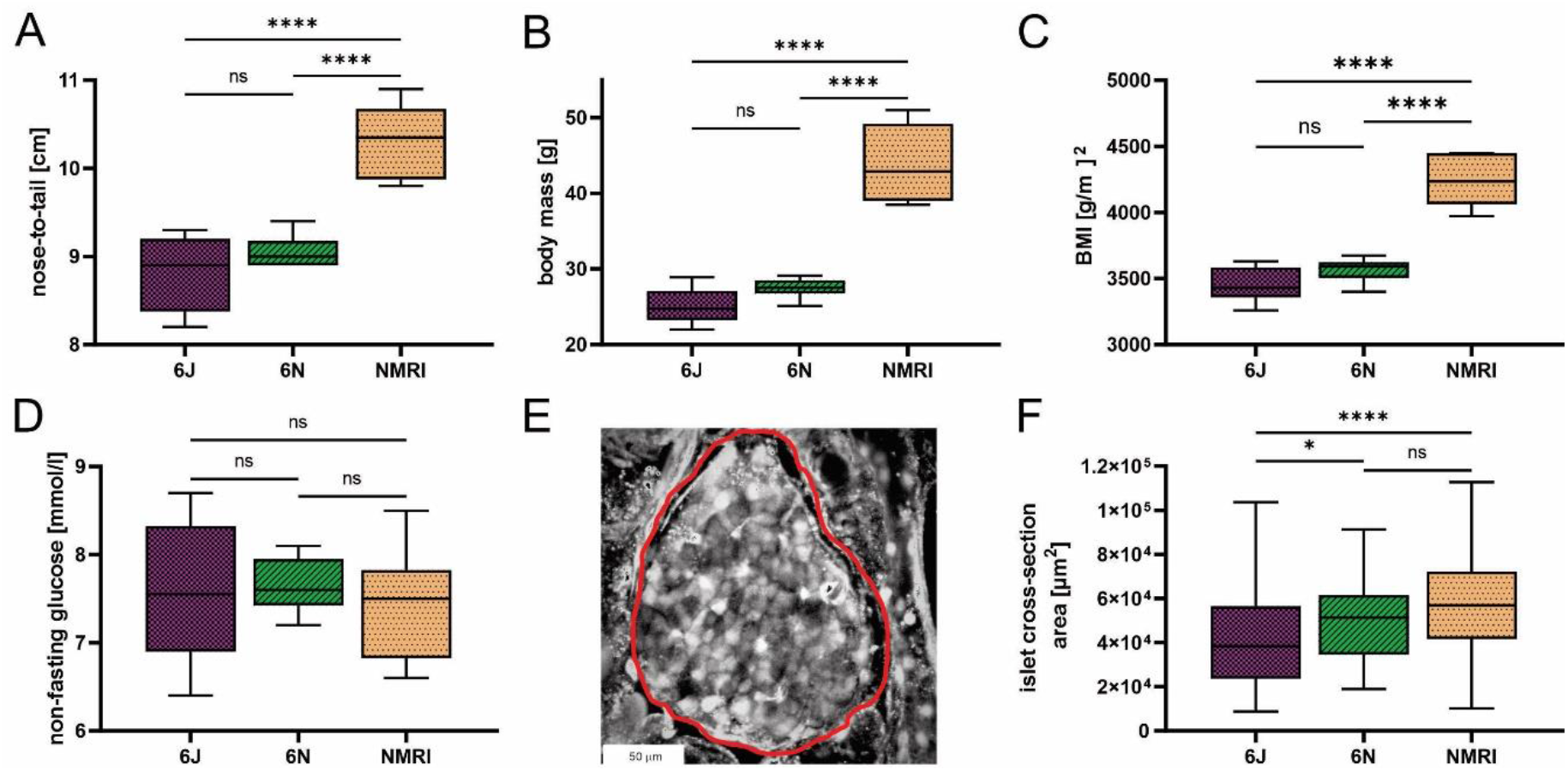
Physiological parameters of C57BL/6J, C57BL/6N, and NMRI mice. **A)** Nose-to-tail length. **B)** Body mass. **C)** Body mass index calculated as the ratio between body weight and body surface area (g/m^2^). **D)** Non-fasting glucose concentration. Data for panels A-D were obtained from 8 C57BL/6J, 6 C57BL/6N, and 6 NMRI mice. **E)** Representative islet of Langerhans. The islet border is marked with a red line. **F)** Islet cross-section area in the pancreas tissue slice (median values 38 373 μm^2^ vs. 51 267 μm^2^ vs. 56 861 μm^2^ for C57BL/6J, C57BL/6N, and NMRI, respectively) obtained from 63/7, 64/6, 75/6 islets/mice from C57BL/6J, C57BL/6N, and NMRI mice respectively.

### Calcium response to glucose stimulation in the three mouse strains

To assess the glucose-dependent properties of activation, plateau phase, and deactivation, we exposed the pancreas tissue slices of the three mouse strains to square-like pulses of stimulatory glucose in the physiological (7, 8, 9, 10 mM) and supraphysiological (12, 16 mM) ranges. A single stimulation was performed per tissue slice. The [Ca^2+^]_i_ dynamics were recorded in beta cells within the optical cross-section of the islet of Langerhans and showed a three-phase response consisting of an initial transient increase in [Ca^2+^]_i_ (activation), followed by a sustained increase with fast oscillations (plateau phase), and a decrease in [Ca^2+^]_i_ after cessation of stimulation (deactivation), as described previously (85).

### The activation time upon stimulation with glucose is delayed in C57BL/6J mice

Beta cells exhibit a characteristic delay following glucose stimulation, a property that has been shown to depend on glucose concentration in the NMRI strain (83, 85). To identify differences between the three strains, we measured the delays between the onset of stimulation and initial [Ca^2+^]_i_ increase in individual beta cells (Figure 2A). Pooling data from several islets of the three strains demonstrated that the delays depended on the glucose concentration in all strains (Figure 2B). Regardless of strain, a general trend was observed, showing that an increase in glucose concentration decreased the median activation delay and that the threshold concentration of glucose was 7 mM. However, the absolute values differed between the three strains. Specifically, delays at most glucose concentrations were longest in the C57BL/6J mice and the shortest in the NMRI mice at all glucose concentrations. The difference in median delays to activation of C57BL/6J mice were by an average of 52.9 % longer than in NMRI mice (85.6 % 7 mM, 41.1 % 8 mM, 50.8 % 9 mM, 89.7 % 10 mM, 28.4 % 12 mM, and 21.4 % 16 mM). The C57BL/6N strain generally showed an intermediate delay that was shorter than in C57BL/6J and longer than in NMRI mice (except at 7 mM and 8 mM glucose, where no difference was found relative to C57BL/6J in the former case and to NMRI mice in the latter). In relative terms, the difference in median delays to activation of C57BL/6N mice was on average 24.6 % shorter than in the C57BL/6J strain. Still, no consistent ratio or trend was observed when comparing different concentrations (−6.4 % 7 mM, 36.9 % 8 mM, 41.0 % 9 mM, 54.5 % 10 mM, 0.4 % 12 mM, and 20.9 % 16 mM). In an attempt to quantify the above strain differences, we calculated the IC50 value (e.g., concentration, at which the delays shortened to 50 % of maximal value) assuming a 4-parameter logistic model (76) (Supplementary Figure 1 (https://doi.org/10.6084/m9.figshare.18356507)). The IC50 values were 8.1 mM (C57BL/6J), 7.8 mM (C57BL/6N), and 6.9 mM (NMRI), demonstrating that the dose-dependent curves of the two inbred strains were right-shifted by 0.9 - 1.2 mM relative to NMRI mice. In this context, we qualitatively observed that some islets of C57BL/6J mice failed to activate in 7 mM glucose in the given time period. However, we did not quantify this further, and the islets were excluded from the quantitative analysis.

**Figure 2:**
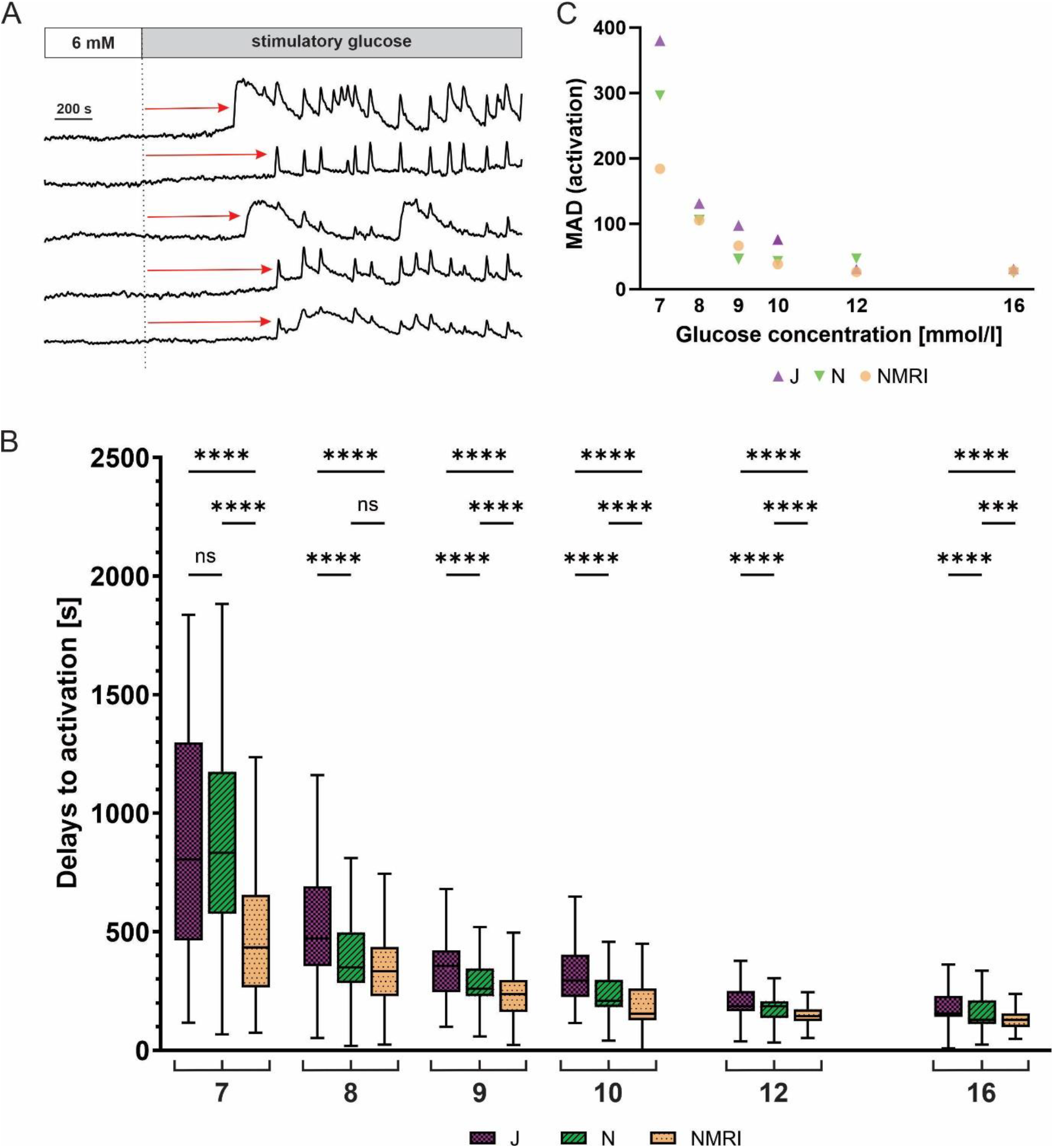
Glucose-dependent activation delays of beta cells in C57BL/6J, C57BL/6N and NMRI mice. **A**) Schematic representation of delays in activation (red arrow) measured as the time of perifusion of the islet with stimulatory glucose (dashed line) before the increase in [Ca^2+^]_i_ signal. **B)** Delays to activation in C57BL/6J (median 805, 472, 357, 294, 186, and 156 s), C57BL/6N (median 833, 349, 260, 210, 185, 129 s), and NMRI (median 434, 334, 236, 155, 145, and 129 s) at 7-, 8-, 9-, 10-, 12-, and 16-mM glucose, respectively. **C)** Variability of activation delays, expressed as median absolute deviation (MAD) after stimulation with 7-, 8-, 9-, 10-, 12-, and 16-mM glucose. MAD values (in seconds): C57BL/6J: 380 (7 mM), 131 (8 mM), 98 (9 mM), 76 (10 mM), 31 (12 mM), and 31 (16 mM), C57BL/6N: 296 (7 mM), 106 (8 mM), 46 (9 mM), 43 (10 mM), 47 (12 mM), and 25 (16 mM), NMRI: 184 (7 mM), 105 (8 mM), 66 (9 mM), 39 (10 mM), 26 (12 mM), and 28 (16 mM). Pooled data (coded as C57BL/6J | C57BL/6N | NMRI) from the following number of cells/islets/pancreas preparations: 239/9/6| 643/13/6| 743/13/6 (7mM glucose), 370/10/7| 876/8/5| 730/12/6 (8 mM glucose), 657/11/7| 851/9/6| 1091/12/6 (9 mM glucose), 521/9/6| 756/9/5| 1078/10/6 (10 mM glucose), 681/11/7| 759/11/6| 904/10/5 (12 mM glucose), and 725/11/5| 703/8/5| 1061/11/6 (16 mM glucose).

We observed a relatively large heterogeneity of activation delays between individual cells of the same islet (Figure 2 A&B). Interquartile ranges were comparable between the inbred and outbred mouse strains. To quantitatively compare the variability among the three strains, we resorted to a robust measure of variability, the median absolute deviation (MAD). Absolute deviations of activation delays from the median were calculated per islet, and the median value of the pooled data was presented in Figure 2C. The heterogeneity of delays was dependent on glucose concentration, as previously reported for the NMRI strain(85). The MAD ranged from the largest values at the threshold concentration (380 s C57BL/6J, 296 s C57BL/6N, 184 s NMRI) to the smallest values at the highest concentration tested (31 s C57BL/6J, 25 s C57BL/6N, 28 s NMRI) (Figure 2C). Excluding the threshold glucose concentration, MAD did not differ substantially between inbred and outbred strains.

### Increasing glucose concentrations modulate the plateau activity differently in the three strains of mice

Following initial activation, beta cells exhibit repetitive oscillations of [Ca^2+^]_i_ that are superimposed on an elevated level of [Ca^2+^]_i_, a feature referred to as the plateau phase of the glucose response (36, 84). To look for possible differences between the three strains during the plateau phase, we measured the duration and frequency of oscillations, as well as the active time, which indicates the proportion of time during which [Ca^2+^]_i_ is elevated (Figure 3A).

**Figure 3:**
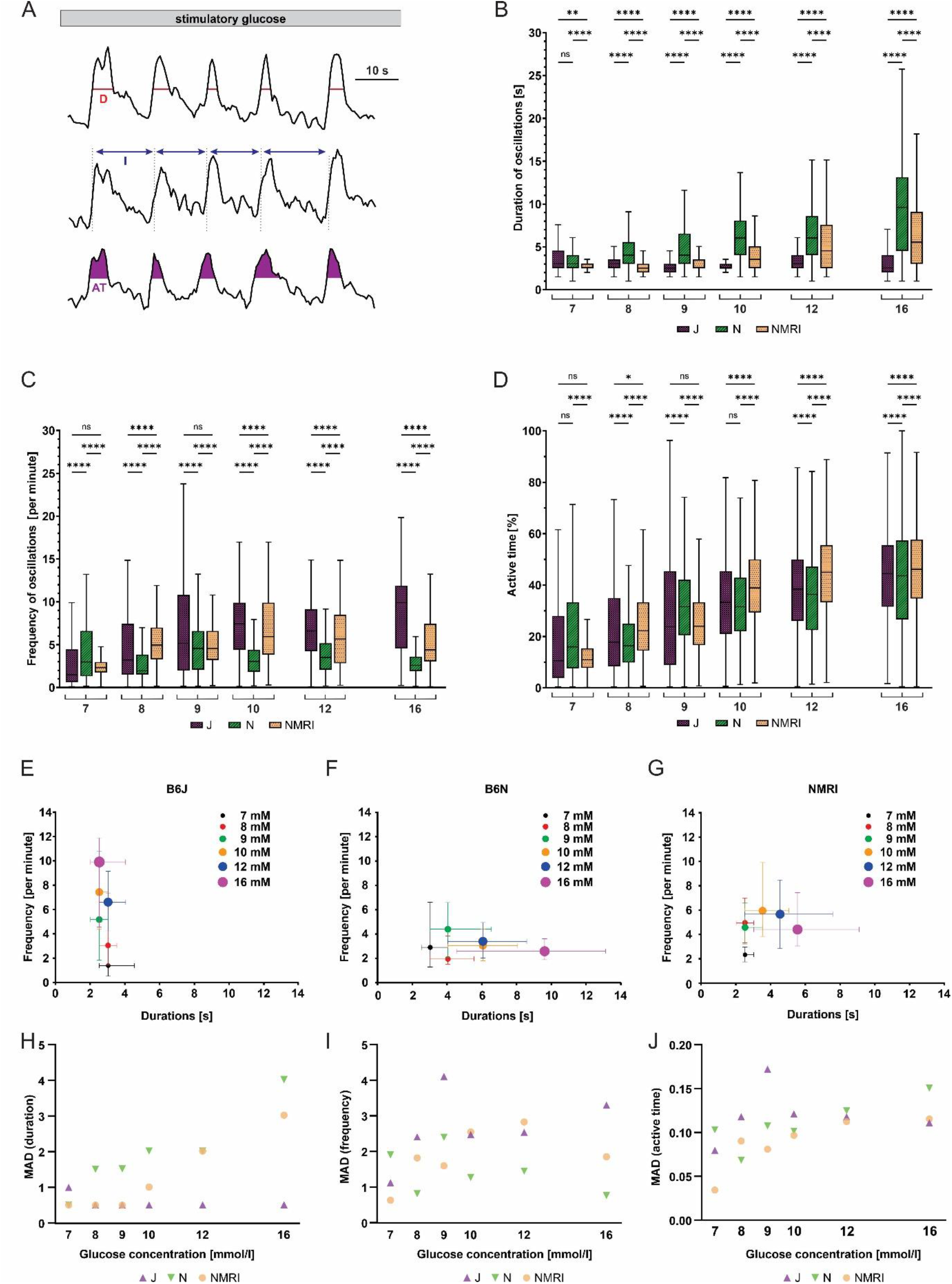
Duration, frequency, and active time of fast oscillations during the plateau phase of beta cell response to glucose in C57BL/6J, C57BL/6N, and NMRI mice. **A)** Schematic representation of analyzed parameters: oscillation duration (D), frequency calculated from burst period (I), and active time (AT). Shown are three typical beta cells during the plateau phase of a response to 10 mM glucose. **B)** Duration of oscillations in the three strains. C57BL/6J: median 3.0, 3.0, 2.5, 2.5, 3.0, and 2.5 s, C57BL/6N: 3.0, 4.0, 4.0, 6.1, 6.1, and 9.6 s, NMRI 2.5, 2.5, 2.5, 3.5, 4.5, and 5.5 s) in 7-, 8-, 9-, 10-, 12-, and 16-mM glucose, respectively. **C)** Frequency of oscillations. C57BL/6J: median (1.5, 3.2, 5.2, 7.4, 6.6, and 9.6 min^−1^, C57BL/6N: 3.0, 1.9, 4.6, 3.0, 3.5, and 2.6 min^−1^, and NMRI: 2.3, 5.0, 4.6, 5.9, 5.7, and 4.4 min^−1^ in 7-, 8-, 9-, 10-, 12-, and 16-mM glucose, respectively. **D)** Percentage of active time. C57BL/6J: median 10.6, 17.8, 23.8, 33.3, 38.5, and 44.4 %, C57BL/6N: median 16.0, 16.4, 31.6, 31.6, 36.4, and 43.6 %, and NMRI: median 10.9, 22.2, 24.0, 38.9, 45.0, and 46.2 % in 7-, 8-, 9-, 10-, 12-, and 16-mM glucose, respectively. **E)** Oscillation durations as a function of respective frequency for B6J. **F)** Oscillation durations as a function of respective frequency for B6N. **G)** Oscillation durations as a function of respective frequency for NMRI. **H)** MAD of oscillation durations. C57BL/6J: median 1.0, 0.5, 0.5, 0.5, 0.5, and 0.5 s, C57BL/6N: median 0.5, 1.5, 1.5, 2.0, 2.0, and 4.0 s, NMRI: 0.5, 0.5, 0.5, 1.0, 2.0, and 3.0 s in 7-, 8-, 9-, 10-, 12-, and 16-mM glucose, respectively. **I)** MAD of oscillation frequency. C57BL/6J: median 1.1, 2.4, 4.1, 2.5, 2.5, and 3.3 min^−1^, C57BL/6N: median 1.9, 0.8, 2.4, 1.3, 1.4, and 0.8 min^−1^, and NMRI: median 0.6, 1.8, 1.6, 2.5, 2.8, and 1.9 min^−1^ in 7-, 8-, 9-, 10-, 12-, and 16-mM glucose, respectively. **J)** MAD of percent of active time. C57BL/6J: median 8.0, 11.8, 17.2, 12.1, 11.8, and 11.1 %, C57BL/6N: 10.3, 6.8, 10.7, 10.2, 12.5, and 15.1 %, and NMRI: median 3.4, 9.0, 8.1, 9.7, 11.2, and 11.5 in 7-, 8-, 9-, 10-, 12-, and 16-mM glucose, respectively. Pooled data (coded as C57BL/6J | C57BL/6N | NMRI) from the following number of cells/islets/pancreas preparations: 57/6/5| 325/11/6| 292/11/6 (7 mM glucose), 96/10/7| 671/8/5 381/10/5 (8 mM glucose), 246/8/7| 684/9/6| 574/10/6 (9 mM glucose), 250/8/5| 531/9/5| 677/9/6 (10 mM glucose), 322/10/7| 456/9/4| 310/7/5 (12 mM glucose), and 256/9/5| 414/7/5| 458/8/6 (16 mM glucose).

Remarkably, when comparing the three strains of mice, we found profound differences in the duration and frequency changes when expressed as a function of the glucose concentration used for stimulation (Figure 3 B&C). More specifically, beta cells from C57BL/6J mice responded to increasing glucose concentrations with an increase in oscillation frequency ranging from a median of 1.5 min^−1^ at 7 mM to 9.9 min^−1^ at 16 mM glucose. In stark contrast, the duration of oscillations remained range-bound to durations like those observed at the threshold concentration (median 2.5 - 3.0 s). The C57BL/6N strain, on the other hand, showed an inverse effect of increasing concentrations: a gradual increase in duration of oscillations rather than their frequency. More specifically, the duration increased from a median of 3.0 s at 7 mM to 9.6 s at 16 mM glucose, while the frequency varied between a median of 1.9 and 4.6 min^−1^, with no clear trend. The NMRI strain, however, modulated active time with frequency in the physiological and with duration in the supraphysiological range of glucose concentration (the shift occurred at glucose concentrations of 9- and 10-mM). Frequency increased from a median of 2.3 min^−1^ at 7 mM to 4.6 - 5.9 min^−1^ at 9 - 10 mM glucose (durations were in the range of median 2.5 and 3.5 s with no clear trend). At glucose concentrations above 10 mM, the duration of oscillations increased to a median of 5.5 s, while the frequency of oscillations even decreased to 4.4 min^−1^ at 16 mM glucose. However, active time increased consistently by an average of 5.4 %, 4.0 %, and 6.3 % per mM increase in glucose in C57BL/6J, C57BL/6N, and NMRI mice, respectively (Figure 3D). We did not observe a consistent difference in active time between the strains. The percentage of active time averaged 12.5 % (C57BL/6J: 10.6 %, C57BL/6N 16.0 %, and NMRI: 10.9 %) at 7 mM glucose and increased to an average of 44.7 % (C57BL/6J: 44.4 %, C57BL/6N 43.6 %, and NMRI: 46.2 %) at 16 mM glucose. To visualize the effect of glucose concentration on frequency and duration, we plotted the frequency of each oscillation against its respective duration, shown in Figure 3 E-G. This approach separated the three strains with respect to the coding of the glucose-stimulated [Ca^2+^]_i_ increase. The C57BL/6J strain depended on the change in frequency to increase the active time (Figure 3E). In contrast, stimulus strength predominately modulated the duration of oscillations in C57BL/6N mice (Figure 3F). The NMRI strain shared properties of both (Figure 3G): The active time depended on an increase in frequency at ≤ 10 mM and the duration at > 10 mM glucose.

Finally, the interquartile range demonstrated a relatively large variability in all three parameters (Figure 3 B-D). However, the variability seemed to change with glucose concentration when the parameter was also concentration-dependent. More specifically, MAD of durations increased with glucose concentration for C57BL/6N (as did the absolute values) and ranged from a median MAD of 0.5 s at 7 mM to 4.0 s at 16 mM glucose, whereas MAD did not change for C57BL/6J (MAD ranged from 0.5 – 1.0 s with no clear trend). Conversely, MAD of frequencies increased in C57BL/6J (as did the absolute values) and ranged from 1.1 min^−1^ at 7 mM to 3.3 min^−1^ at 16 mM glucose, while the duration remained unaffected by glucose concentration (median MAD 0.8 – 2.4 min^−1^ with no clear trend). The NMRI strain exhibited both properties, increasing MAD of frequencies in the 7 – 10 mM glucose range (MAD 0.6 min^−1^ at 7 mM and 2.5 min^−1^ at 16 mM glucose), while MAD of durations increased above 10 mM glucose (MAD duration 3.0 s at 16 mM glucose). MAD of active time increased with increasing glucose concentrations in all strains spanning from 8.0, 10.3, and 3.5 % in 7 mM glucose to 11.1, 15.1, and 11.5 % in 16 mM for C57BL/6J, C57BL/6N, and NMRI mice. No systematic differences were observed in MAD of active time between inbred and outbred mice.

### C57BL/6J mice deactivate earlier following stimulation

Following glucose withdrawal, beta cells cease their oscillatory activity and return to the pre-stimulatory baseline [Ca^2+^]_i_ after a certain time delay (Figure 4A), as demonstrated previously for the NMRI strain (83, 85). To search for differences in the deactivation properties, we measured the deactivation delays of individual cells in all three strains. Generally, all three strains showed a concentration-dependent lengthening of deactivation delays (Figure 4B). However, in experiments using physiological glucose concentrations (7 - 10 mM glucose), an intermittent period of no activity in a portion of cells preceded or coincided with the withdrawal of the stimulus. The rest of the cells subsided their activity during and after stimulus withdrawal. The intermittent inactivity at the time of stimulus withdrawal resulted in a portion of negative deactivation delays. This effect was most noticeable in the C57BL/6J strain of mice, where the effect subsided glucose-dependently from 7 mM to 9 mM glucose and was negligible in higher (≥ 10 mM) glucose concentrations. Regardless of this phenomenon, increasing deactivation delays were observed, ranging from −208 s at 7 mM to 370 s at 16 mM glucose. The negative deactivation delays were also seen in C57BL/6N and NMRI mice, but much more rarely and mainly at concentrations of glucose < 8 mM. Similarly, for these two strains, the deactivation delays were glucose-dependent, ranging from a median of 133 s (C57BL/6N) and 178 s (NMRI) at 7 mM to 386 s (C57BL/6N) and 278 s (NMRI) at 16 mM glucose, respectively. Delays to deactivation were typically the longest in NMRI mice and the shortest in C57BL/6J mice. On average, the difference in median delays between NMRI and C57BL/6J mice was 47.1 % (216.8 % for 7 mM, 27.8 % for 8 mM, 6.3 % for 9 mM, 15.7 % for 10 mM, 13.8 % for 12 mM, and 2.3 % for 16 mM). The C57BL/6N mice presented with an irregular deactivation pattern compared to C57BL/6J and NMRI mice, being the first to deactivate at some glucose concentrations (9 and 12 mM) and last in others (8 and 16 mM). However, their deactivation times more closely resembled those of NMRI mice than C57BL/6J mice. More specifically, the average difference in median delays between NMRI and C57BL/6N mice was 9.9 % (36.5 % for 7 mM, −17.9 % for 8 mM, 9.4 % for 9 mM, 7.9 % for 10 mM, 25.8 % for 12 mM, and −2.0 % for 16 mM).

**Figure 4:**
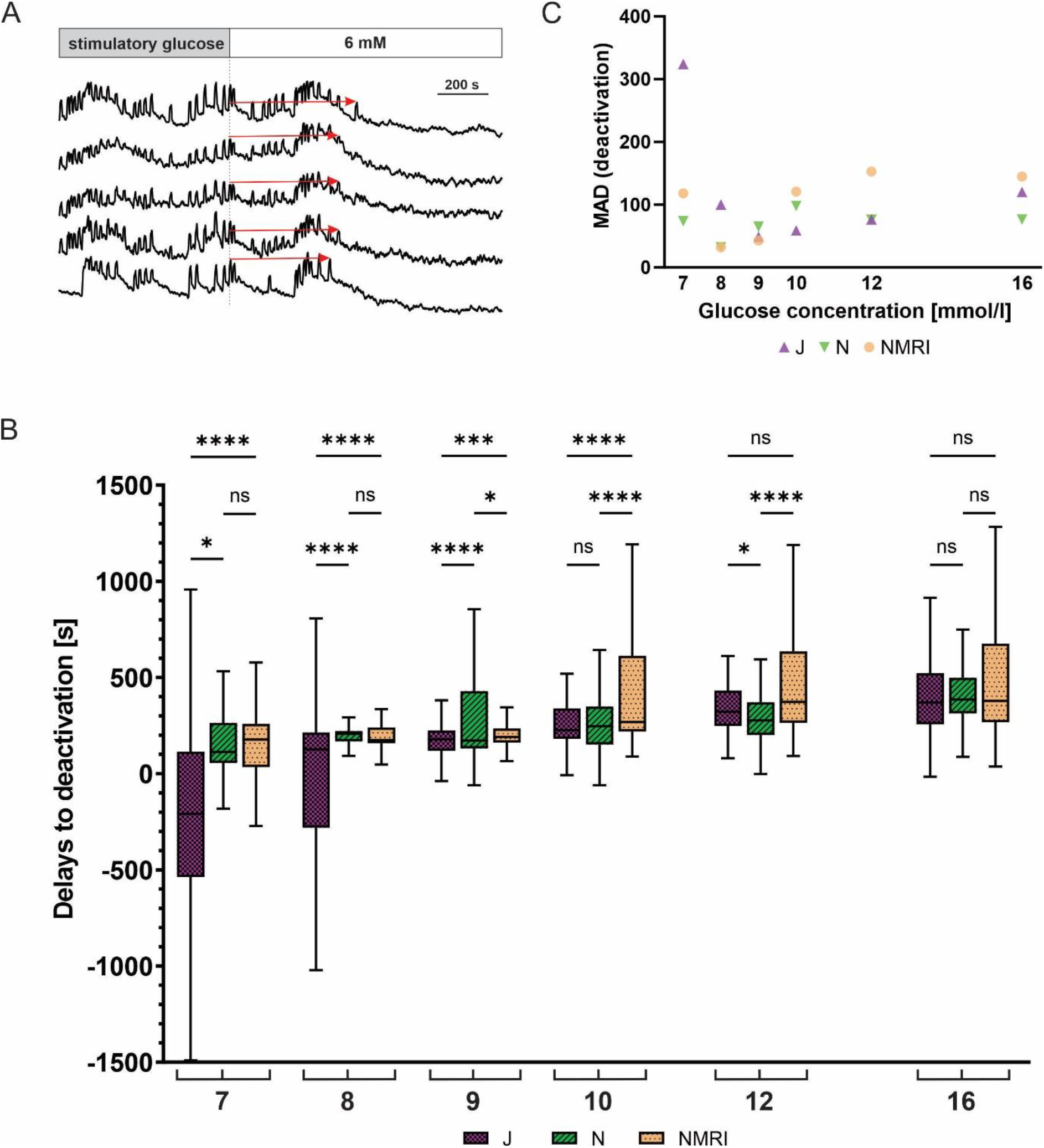
Glucose-dependent deactivation delays of beta cells in C57BL/6J, C57BL/6N and NMRI strains. **A)** Schematic representation of delays to deactivation (red arrow) measured as the time difference between end of stimulation (dashed line) and the last [Ca^2+^]_i_ oscillation. **B)** Delays to deactivation in C57BL/6J (median −208, 126, 179, 227, 322, and 370 s), C57BL/6N (median 113, 206, 173, 248, 277, and 386 s), and NMRI (median 178, 175, 190, 269, 373, and 378 s) in 7-, 8-, 9-, 10-, 12-, and 16-mM glucose, respectively. **C)** Variability in deactivation delays, expressed as median absolute deviation (MAD) after stimulation with 7-, 8-, 9-, 10-, 12-, and 16-mM glucose. MAD values (in seconds): C57BL/6J: 324 (7 mM), 100 (8 mM), 49 (9 mM), 59 (10 mM), 77 (12 mM), and 120 (16 mM), C57BL/6N: 74 (7 mM), 32 (8 mM), 65 (9 mM), 98 (10 mM), 76 (12 mM), and 77 (16 mM), NMRI: 118 (7 mM), 32 (8 mM), 43 (9 mM), 121 (10 mM), 153 (12 mM), and 145 (16 mM). Pooled data (coded as C57BL/6J | C57BL/6N | NMRI) from the following number of cells/islets/pancreas preparations: 133/6/5| 377/12/6| 472/12/6 (7mM glucose), 288/10/7| 832/8/5| 482/11/6 (8 mM glucose), 544/10/7| 700/9/5| 792/10/6 (9 mM glucose), 307/5/4| 711/9/5| 999/10/6 (10 mM glucose), 546/10/6| 676/11/6| 749/10/6 (12 mM glucose), and 657/11/6| 592/7/5| 809/10/6 (16 mM glucose).

Similarly to the activation, we observed relatively large variability in the deactivation delays (Figure 4C). We calculated the MAD of deactivation delays of cells in an islet and plotted pooled median values to display interstrain differences. Regarding the glucose concentration, the variability in the C57BL/6J expressed a U-shape with peaks observed at 7 mM (324 s) and 16 mM (170 s) glucose and with a nadir at 9 mM (49 s). The C57BL/6N and NMRI similarly exhibited a U-shaped dose-dependent response (C57BL/6N: peaks 74 s (7 mM) and 98 s (10 mM), NMRI: peaks 118 s (7 mM) and 153 s (12 mM)) that saturated (and even decreased) at higher stimulatory glucose concentrations. As with activations, no clear differences were observed between outbred and inbred strains.

## DISCUSSION

Although animal models are generally accepted in (patho)physiological research of the pancreas, study outcomes were often generalized across taxonomic genera or even orders, assuming the same phenotypes and (patho)physiological mechanisms. This was an unsound assumption even for the most commonly used animal model *Mus musculus* (laboratory mouse), as several studies revealed significant differences between strains, especially in terms of animal size (72), glucose metabolism (5, 55), and even susceptibility to disease (21, 42, 58, 68), to name a few. In the present study, we attempted to at least partly overcome this problem by systematically comparing the beta cell responses to glucose in the three most commonly used mouse models, the outbred NNT competent NMRI mice and the two inbred C57BL/6 substrains, i.e., the NNT deficient C57BL/6J and the NNT competent C57BL/6N mice. Apart from our own previous work (85), which was limited to the NMRI strain, to the best of our knowledge, the glucose-dependency of activation, activity, and deactivation of beta cells in intact islets of Langerhans has not been systematically studied.

To robustly assess how the analyzed beta cell characteristics relate to the *in vivo* glycemic status in these three strains, we also examined some basic physiological parameters. NMRI mice were larger and heavier than either of the C57BL/6 substrain, consistent with previous findings (62). Unsurprisingly, the same tendency was observed for the calculated BMI. Non-fasting glucose did not differ between the strains and was in accordance with previous results (49). The larger cross-sectional area of the islets in the NMRI strain may be due to a greater insulin requirement on account of larger animal size and contribute to normal glucose tolerance in these animals, given the very similar active time responses. Islet number clearly increases with body size across different species, but a possible contribution of islet size to increasing total islet mass in larger organisms is much less clear (15, 22, 43, 44, 56). It has been reported that at least within species, islet size and architecture can change with increased demand due to growth and pregnancy(14, 56, 69). Islet areas of C57BL/6N and NMRI mice have not yet been systematically compared and our findings offer some support to the idea that increased islet size may account for at least a part of increased total islet mass in larger organisms. However, our cross-sectional data from a few slices are subject to many caveats. The most important is a possible bias due to the selection of islets for calcium imaging. Future studies aimed at either random sampling of islet cross-sections within tissue slices, combined with insulin secretion data from the same slices, could provide more insight. However, it is worth mentioning that judging by previous work, the number of islets in NMRI and C57BL/6 mice of similar age probably does not differ significantly (14, 69).

Beta cells respond to a glucose stimulus with a time delay, a phenomenon attributed to the time needed for beta cells to metabolize glucose. Previously, large differences between individual cells following a random activation pattern have been reported for the NMRI strain (83, 85). The present results confirm the previous findings in NMRI mice on an independent sample and further show that there is similar activation heterogeneity at the level of individual cells in all three substrains (Figure 2B). Importantly, beta cells show a progressive decrease in delays with increasing glucose concentration, a phenomenon termed advancement of beta cell activation, as previously demonstrated for the NMRI strain (85). The advancement of activation was observed in all three studied mouse strains (Figure 2B). The delay effect displayed a minimum at approximately 2 min, which agrees well with results observed in rats (70). However, the concentration dependence did not overlap when the three mouse strains were compared. Both inbred strains were right-shifted by 1.2 (C57BL/6J) and 0.9 mM (C57BL/6N) compared to NMRI mice (Supplementary Figure 1 (https://doi.org/10.6084/m9.figshare.18356507)), suggesting that a higher glucose concentration is required for a comparable initiation of activation. Delays in activation were most pronounced in C57BL/6J mice resulting in several islets of C57BL/6J mice failing to respond to the established threshold for stimulatory glucose (7 mM). Although a comparable threshold concentration of 7.1 mM (Figure 2B) has been reported for insulin release from isolated islets of fed and food-deprived C57BL/6J mice (18), a failure to report unresponsive islets, due to a biased search for responsive islets could have influence concentration-dependence curves. The discrepancy in threshold concentration between our and previous results could also be attributed to differences in pre-stimulatory glucose concentration (71), a broadening of the curve of concentration dependence as a consequence of islet preparation, or other factors(46, 53).

After the initial activation described above, beta cells exhibit a plateau phase of [Ca^2+^]_i_ activity characterized by either fast, slow, or mixed oscillations (11, 12, 89, 94). How beta cells encode glucose concentration during this plateau phase is still a matter of debate. In tissue slices, fast oscillations are typically observed and previous studies that focused on the fast component provided conflicting results. Some showed an increase in the duration of individual oscillations (6), while others reported increased frequency as glucose concentration (57) increased. Many causes for this discrepancy are conceivable: sample preparation (damage due to enzymatic isolation of cells and islets, mechanical disruption due to microdissection of islets, lack of neuronal innervation in the tissue slice approach, poorly defined conditions for *in vivo* measurements), temporal resolution (high in electrophysiological recordings, low during confocal imaging), spatial resolution (single cell resolution when using patch pipette, islet resolution when using photomultiplier, multicellular resolution when using confocal imaging), the hormonal status of the animal (hormonal cycles in females), and inter-species and - strain differences, to name only a few. In this study, we compared plateau phases of beta cells in three strains of mice to account for the inter-strain effect. Surprisingly, the three strains of mice could be divided into three different categories, each with its own characteristic coding principles of either tuning the frequency, the duration, or a combination of both (Figure 3). Beta cells from C57BL/6N mice exclusively modulated the duration, which tripled over the tested range (from ~ 3 s at 7 mM to ~ 10 s at 16 mM glucose), while the frequency remained unchanged (~ 3-4 min^−1^). The C57BL/6J mice showed almost exactly the opposite characteristics: an exclusive modulation of oscillation frequency that increased approximately threefold over the tested range (from ~ 1.5 - 3 to ~ 9 min^−1^). The NMRI strain exhibited the most complex coding strategy, switching from increases in frequency to increases in duration with increasing levels of stimulation. At 7 – 10 mM, the frequency increased approximately threefold (from ~ 2 in the lower to ~ 6 min^−1^ in the upper range), while durations remained unchanged. In yet higher glucose, durations increased about twofold (from ~ 2.5 s to ~ 5.5 s at 16 mM glucose), while frequency remained unchanged. To graphically represent the three coding modes, we plotted frequency as a function of respective durations over the tested range of concentrations (Figure 3 E-G). This approach clearly separated the three substrains in terms of how the two parameters interacted: (i) exclusive frequency modulation in C57BL/6J mice, (ii) exclusive duration modulation in C57BL/6N mice, and (iii) a combination of both in NMRI mice. To our knowledge, this is the first report of such pronounced differences between substrains of the same species. The different properties should be carefully considered when interpreting the effects of various physiological or pharmacological secretagogues on oscillations to not miss a possible effect on frequency when there is no effect on duration, and when comparing the results from different studies to avoid the erroneous assumption that “a mouse is a mouse”.

Surprisingly, the three strains shared remarkably similar curves of active time. Active time is best understood as the fraction of time that the calcium signal is elevated during repetitive oscillations. The values of ~ 15 % at the threshold concentration increased to ~ 45 % at 16 mM in an almost linear fashion in all substrains. In other words, for an increase in glucose concentration by 9 mM over the threshold level of 6 mM, the active time increased by 30 %, or approximately 3 % per 1 mM glucose.

It is unclear how the increase in active time in all three mouse strains translates into effects on insulin secretion, per se, as only fast [Ca^2+^]_i_ oscillations were considered in this study. For the NMRI, we have recently shown that the active time of the fast oscillatory component almost perfectly followed the insulin secretion data (85). In contrast, the slow component was independent of glucose concentration (94). However, beta cells can most likely increase their active time to an even higher proportion if stimulated with even higher glucose concentrations than that used in this study. For instance, electrical activity of beta cells (24) was shown to approach saturation at 20 mM glucose or even higher and similar findings have been shown for insulin secretion (2, 8, 10, 32, 47, 60). In fact, insulin secretion appears to increase well beyond what has been previously reported via [Ca^2+^]_i_-independent mechanisms (34). Considering active time increases similarly in all three mouse strains, but glucose-stimulated insulin release is impaired in C57BL/6J mice (55, 87), insulin secretion could be impaired not only upstream but also downstream of calcium at the level of the exocytotic machinery. This is not yet clear, as there are findings in favor of the former (20) and the latter (30, 87). In future studies, it would be prudent to assess active time (and other oscillation parameters) also in 20 mM or higher glucose and also determine into more detail the relationship between active time estimated from electrical recordings and calcium oscillations at half-amplitude. The latter could theoretically yield lower values than the former and this discrepancy may account for slightly lower values of active time in this study compared with some previous findings (45, 63).

Finally, the deactivation properties of beta cells at the level of [Ca^2+^]_i_ have not yet been systematically examined for differences between strains. In all three strains, beta cell oscillatory [Ca^2+^]_i_ activity decreases when the stimulatory glucose is removed. However, it should be noted that in some islets, certain cells remained active to a lesser extent even 20 – 40 min after the end of stimulation, coinciding with an initial pause in [Ca^2+^]_i_ activity. This was observed mainly in the C57BL/6N and the NMRI mice, mostly at physiological glucose concentrations. However, quantification of these results could not be performed because it was impossible to establish strict criteria for categorizing these cells due to the varying length of the individual recordings. Therefore, this information was not included in our analysis. As for the cells that deactivated, data from all three strains of mice showed that the time required for deactivation increased with increasing glucose concentration. In some islets of the C57BL/6J strain, oscillations ceased even before the non-stimulatory glucose was reintroduced, resulting in negative delays in activation. This occurred at 7 – 9 mM stimulatory glucose concentrations, again indicating a shifted threshold for glucose-induced insulin secretion in the C57BL/6J strain. When stimulatory glucose was increased, delays in deactivation were prolonged and similar in all three groups.

The variability in activations, expressed by the median absolute deviation from the median, showed that variability was the highest at lower glucose concentrations, indicating that activations are much more dependent on activation thresholds within individual cells. As stimulatory glucose concentration increased, the variability of delays decreased, suggesting that activation of cells within an islet at supraphysiological concentrations is more strongly tied to the activity of other cells. In the plateau phase, variability increased with increasing glucose concentrations for the NMRI and C57BL/6N strains as far as durations are concerned but increased up to 12 mM glucose and decreased at 16 mM when looking at the variability of frequency. In C57BL/6J mice, variability stayed fairly constant in the case of durations, but the variability of oscillation frequencies increased steadily. Variability in active time increased with higher glucose concentrations for all strains. Variability in deactivations was higher at the threshold and in supraphysiological concentrations. We surprisingly did not observe any systematically higher variability in the outbred NMRI mice compared to the C57BL/6 substrains for any of the assessed parameters. This is somewhat at odds with recent blanket statement appeals for preferential use of inbred mouse strains over outbred stocks in research (27) and deserves further investigations in the future.

## CONCLUSION

In this study, we characterized for the first time glucose-stimulated [Ca^2+^]_i_ dynamics in two inbred mouse strains (C57BL/6J and C57BL/6N) most commonly used in islet research and confirmed our previous findings in outbred NMRI mice on a new independent sample (83, 85). We wish to point out that the use of inbred mice doesn’t necessarily reduce variability, which is particularly true for experiments at physiological glucose concentrations. In future studies, findings on [Ca^2+^]_i_ dynamics could be correlated with electrophysiological and insulin secretion data (something not yet possible in conjunction with the tissue slice). In addition, it will be interesting to see whether the [Ca^2+^]_i_ waves in the three substrains are comparable and what their functional connectivity patterns are (81), together with possible changes in dietary and genetic models of diabetes based on different backgrounds. Finally, our findings also underline the general importance of transparent experimental design, analysis, and interpretation of results (4). The apparent phenotypic differences between beta cells even within the same species should be accounted for when comparing the results of different studies and when examining the differences between mouse islets and islets of other species, since a mouse is not a mouse. Such an approach will hopefully help reconcile some of the currently conflicting views about beta cell physiology in mice, lead to a more accurate translation to humans, and ultimately contribute to a better understanding and treatment of diabetes.

## Acknowledgement

We thank Maruša Rošer, Rudi Mlakar, and Eva Paradiž Leitgeb, for their excellent support.

## Author contribution

**Conceptualization:** Andraž Stožer, Jurij Dolenšek,

**Formal analysis:** Viljem Pohorec, Jurij Dolenšek, Andraž Stožer, Maša Skelin Klemen, Lidija Križančić-Bombek

**Software:** Jurij Dolenšek,

**Experimental work:** Viljem Pohorec, Lidija Križančić-Bombek, Andraž Stožer, Maša Skelin Klemen, Jurij Dolenšek

**Funding acquisition:** Andraž Stožer,

**Project administration:** Andraž Stožer,

**Supervision:** Andraž Stožer, Jurij Dolenšek

**Visualization:** Maša Skelin Klemen, Jurij Dolenšek, Andraž Stožer

**Writing – original draft:** Viljem Pohorec, Jurij Dolenšek, Andraž Stožer,

**Writing – review & editing:** Viljem Pohorec, Jurij Dolenšek, Andraž Stožer, Maša Skelin Klemen, Lidija Križančić-Bombek,

