## Supplemental Figure 1 for "Glucose-stimulated calcium dynamics in beta cells from C57BL/6J, C57BL/6N, and NMRI mice: A systematic comparison of activation, activity, and deactivation properties in tissue slices"

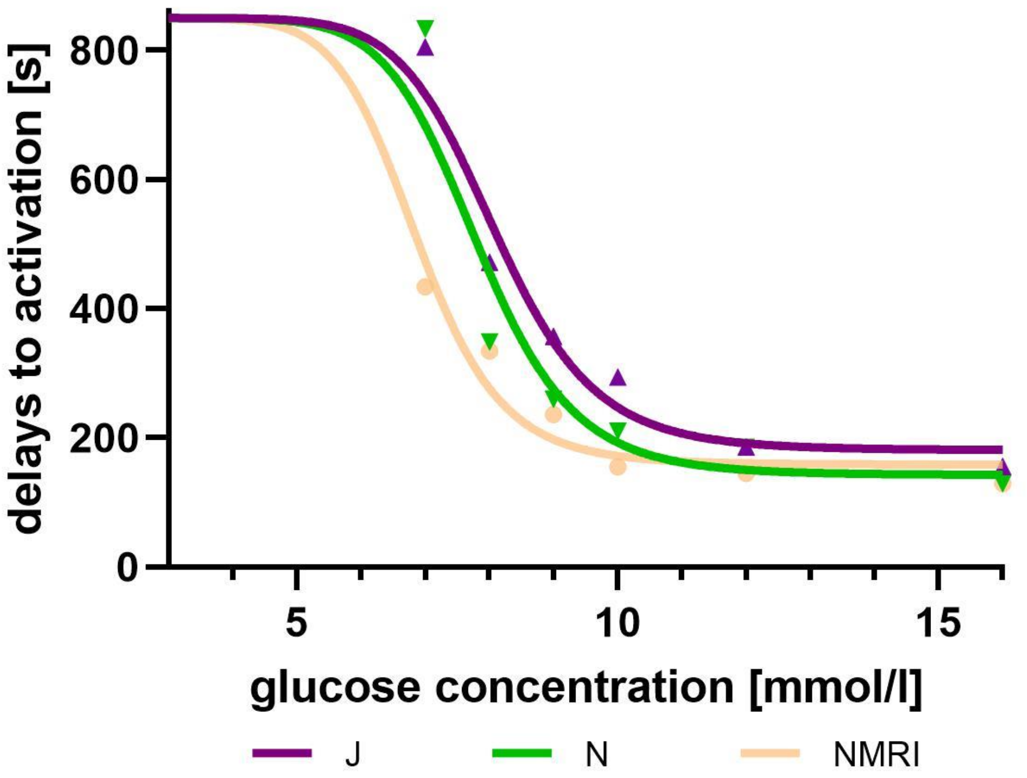


**Supplementary Figure 1: Activation delays of beta cells are left-shifted in NMRI mice compared to the C57BL/6J and C57BL/6N.** Fitted 4-parameter logistic regression model of delays in activation of beta cells in C57BL/6J, C57BL/6N and NMRI mice at 7-, 8-, 9-, 10-, 12-, and 16- mM glucose. Delays to activation in C57BL/6J (median 805, 472, 357, 294, 186, and 156 s), C57BL/6N (median 833, 349, 260, 210, 185, 129 s), and NMRI (median 434, 334, 236, 155, 145, and 129 s) at 7-, 8-, 9-, 10-, 12-, and 16- mM glucose, respectively. Pooled data (coded as C57BL/6J | C57BL/6N | NMRI) from the following number of cells/islets/pancreas preparations: 239/9/6| 643/13/6| 743/13/6 (7mM glucose), 370/10/7| 876/8/5| 730/12/6 (8 mM glucose), 657/11/7| 851/9/6| 1091/12/6 (9 mM glucose), 521/9/6| 756/9/5| 1078/10/6 (10 mM glucose), 681/11/7| 759/11/6| 904/10/5 (12 mM glucose), and 725/11/5| 703/8/5| 1061/11/6 (16 mM glucose).
